# Validity of natural isotope abundance correction for metabolic flux analysis

**DOI:** 10.1101/2020.05.04.075838

**Authors:** Roland Nilsson

## Abstract

A pervasive issue in stable isotope tracing and metabolic flux analysis is the presence of naturally occurring isotopes such as ^13^C. For mass isotopomer distributions (MIDs) measured by mass spectrometry, it is common practice to correct for natural isotopes within metabolites of interest using a particular linear transform based on binomial distributions. However, the origin and mathematical derivation of this transform is rather obscure, and it may be difficult for nonexperts to understand precisely how to interpret the resulting corrected MIDs. Moreover, corrected MIDs are often used to fit metabolic network models and infer metabolic fluxes, which implicitly assumes that corrected MIDs will yield the same flux solution as the actual observed MIDs. Yet, there seems to be no published proof of this important property. Here, we provide a detailed derivation of the MID-correcting linear transform, reflecting its historical development, and describe some interesting properties. We also provide a proof that for metabolic flux analysis on noise-free MID data at steady state, observed and corrected MIDs indeed yield the same flux solution. On the other hand, for noisy MID data, the flux solution will generally differ between the two representations.

## 1 Introduction

When analyzing data from stable isotope tracing experiments, the presence of naturally occurring isotopes can be problematic. For example, carbon-13 (^13^C) accounts for about 1% of all carbon in the biosphere [Berglund and Wieser, 2011], and when measuring metabolic products we must somehow correct for this natural ^13^C to determine the amount of ^13^C derived from a tracer [Buescher et al., 2015]. Obviously, there is no physical difference between naturally occurring and tracer-derived ^13^C, so this correction can only be made statistically. To make this notion precise, we will need some definitions. For mass spectrometry data, an *n*-carbon compound can be divided into *n* + 1 mass isotopomers (also called isotopologues), where mass isotopomer *i* represents the set of molecules of the compound that contain exactly *i* ^13^C atoms (*i* = 0, 1, …, *n*), regardless of their position in the molecule. We will consider only ^13^C in this paper, but other isotopes such as ^15^N or ^2^H can be handled similarly. The associated mass isotopomer distribution (MID) is an (*n* + 1)-vector *x* = (*x*_0_ *x*_1_ … *x*_*n*_)^*T*^ where *x*_*i*_ is the fraction of molecules having *i* ^13^C atoms. (Throughout, we will index MID vectors and related matrices starting from zero for convenience.) Because natural ^13^C atoms occur randomly and independently with a probability *p* ≈ 0.01, the “natural” mass isotopomer distribution (MID) of an *n*-carbon compound is a vector *β*^*n*^ whose elements follow the binomial distribution

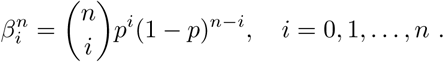

In isotope tracing experiments, natural-derived and tracer-derived ^13^C inevitably mix to form the observed MIDs. An MID correction method is then a function that maps any observed MID *x* to a hypothetical MID *y* that *would have been observed* if natural ^13^C did not exist. We refer to *y* as a *corrected* MID. In particular, MID correction must map the natural MID *β*^*n*^ to the unit vector *e*^*n*,0^ = (1 0 … 0)^*T*^. Over the years, several methods and software packages have been developed for MID correction [Millard et al., 2012, Jungreuthmayer et al., 2015, Su et al., 2017, Millard et al., 2019], providing various features such as support for multiple isotopes and compound fragmentation. They are all based on a particular class of linear transforms *T*^*n*^, *n* = 1, 2, … that relate an *n*-carbon MID *x* to the corrected MID *y* as *x* = *T*^*n*^*y*, where *T*^*n*^ is the (*n* + 1) × (*n* + 1) matrix

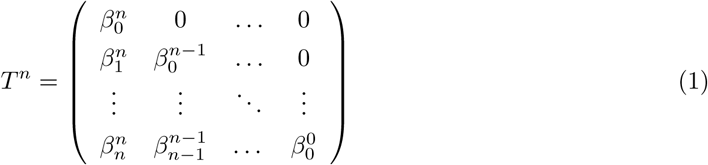

MID correction then simply amounts to solve this system for *y* = (*T*^*n*^)^−1^*x* (the matrix *T*^*n*^ is always invertible). However, it is difficult to find literature explaining how this particular transform is derived and demonstrating why the MID *y* is indeed a corrected MID, as defined above. The transform *T*^*n*^ was developed in a series of technical studies going back as far the 1960’s [Brauman, 1966], but these may be inaccessible to non-experts, and so the origin and underlying logic of the transform *T*^*n*^ remains somewhat obscure. An excellent survey of the historical development of MID correction was given by Midani et al. [2017], but did not include a mathematical derivation of the transform *T*^*n*^. In section 2 of this paper, we provide such a derivation, reflecting the historical development. We then discuss some interesting properties of *T*^*n*^ in section 3.

Another important question arises when the corrected MIDs *y* are used for metabolic flux analysis (MFA). Performing MFA on corrected MIDs is attractive since we need not keep track of natural isotopes, simplifying the analysis [Buescher et al., 2015]. But, cellular metabolism obviously acts on the “uncorrected” MIDs *x* that physically occur, and therefore MFA on these uncorrected MIDs (where “unlabeled” molecules have binomial MIDs *β*^*n*^) must be regarded as the ground truth. Therefore, to ensure that MFA on the corrected MIDs *y* is valid, we must first establish that this always yields the same results as MFA on the actual MIDs *x*. To our knowledge there is no published proof of this proposition. We provide such a proof for the case of noise-free data at isotopic steady-state in section 4, and conclude by discussing some remaining open questions.

## 2 Derivation of the MID correction

To understand the origin of the MID correcting transform *T*^*n*^, we here provide a derivation reflecting its historical development [Midani et al., 2017]. We will first review mixture models of MIDs, which constitute a related but somewhat different type of “correction” that yields the fractional abundance of one or more tracers in a metabolite pool, thereby accounting for natural ^13^C. We then describe a general model for the MID of labeled compounds that occur in mixture models, and finally show how these models can be generalized to obtain the transform *T*^*n*^.

### 2.1 Linear mixtures of tracers

We first consider the simple case where an *n*-carbon tracer with known MID *x*^1^ is mixed with an endogenous source of the same compound, which has natural MID *x*^0^ = *β*^6^ (Figure 1). At steady state, the observed (measured) MID *x* is then a linear mixture

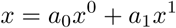

where *a*_0_, *a*_1_ are the mixture fractions (satisfying *a*_0_ + *a*_1_ = 1). This situation typically arises in isotope tracing *in vivo*, where *x*^0^ is known as the *tracee* [Cobelli et al., 1992]. Provided that we know the tracer MID *x*^1^ and can measure the MID *x*, the above yields *n* + 1 equations and two unknowns, and we can solve for the mixture coefficients *a* = (*a*_0_, *a*_1_) using a least-squares method [Brauman, 1966]. For the example data given in Figure 1, this gives *a* = (0.85, 0.15), indicating that the tracer constitutes 15% of the observed mixture, after “correction” for natural ^13^C. As expected, *a*_1_ < *x*_1_ = 0.191, since natural ^13^C also contributes to *x*_1_. It is straightforward to extend this model to multiple tracers with MIDs *x*^1^, *x*^2^, …, *x*^*K*^,

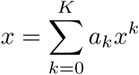

**Figure 1:**
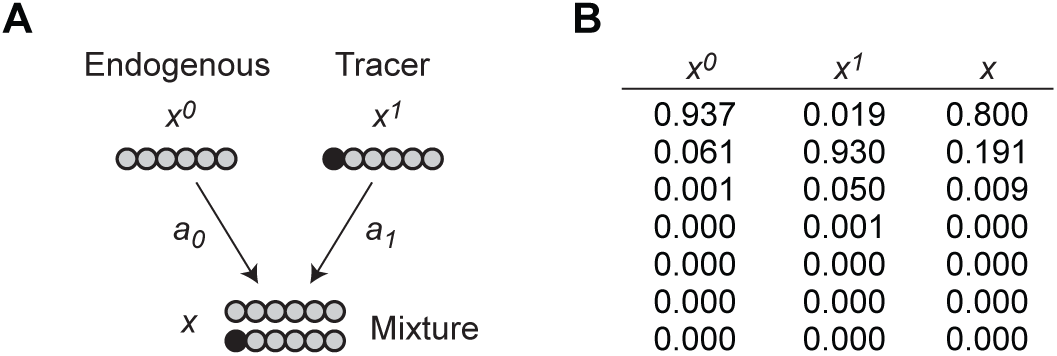
A tracer-tracee example. (A) An endogenous 6-carbon compound with natural MID *x*^0^ = *β*^6^ is mixed with a 1-^13^C tracer *x*^1^ in proportions *a*_0_, *a*_1_, forming a mixture with MID *x*. Black circles indicate ^13^C atoms, gray circles atoms drawn from the natural isotope distribution. (B) Example (hypothetical) MID data from the system in A.

Provided that *K* ≤ *n*, we can still solve for the mixture coefficients *a*_*k*_.

An important generalization is to apply the above mixture model also in cases where some of the mixture components are metabolic products. For example, Rosenblatt et al. [1992] described a case where a doubly labeled leucine tracer is metabolized to a singly labeled one, and both are mixed with endogenous leucine (Figure 2). To solve this model, we must of course know the MID *x*^1^ of the biologically synthesized ^13^C_1_-leucine, as well as the tracer MID *x*^2^. Since *x*^1^ cannot be observed experimentally, Rosenblatt et al. [1992] suggested to instead use a theoretical distribution, which we will describe next.

**Figure 2:**
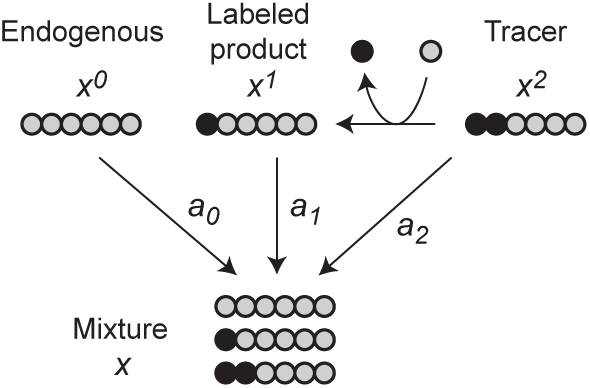
Mixture model adapted from Rosenblatt et al. [1992], with some simplifications. Here a ^13^C_2_ leucine tracer with MID *x*^2^ is metabolized to a ^13^C_1_-leucine compound (*x*^1^), and both are mixed with endogenous leucine (*x*^0^), with mixture coefficients *a*_*k*_.

### 2.2 The MID of labeled compounds

In this section, we describe the MID of an *n*-carbon compound that has incorporated *k* atoms from a pure ^13^C source and *n* − *k* atoms from a natural distribution, 0 ≤ *k* ≤ *n*. Denote this MID *τ*^*n,k*^. For the special case *k* = 0 we simply have *τ*^*n*,0^ = *β*^*n*^, while for *k* = *n* we have the pure ^13^C_*n*_ MID *τ*^*n,n*^ = *e*^*n,n*^. For 0 < *k* < *n*, we assume that the *n*-carbon compound is formed by a condensation between a pure ^13^C_*k*_ compound with MID *τ*^*k,k*^ and a naturally distributed compound with MID *τ*^*n*−*k*,0^ (see Figure 3A for an example). The resulting MID *τ*^*n,k*^ of the condensation product is then given by the convolution

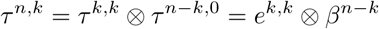

where *x* ⊗ *y* is defined component-wise as

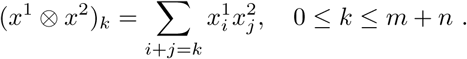

**Figure 3:**
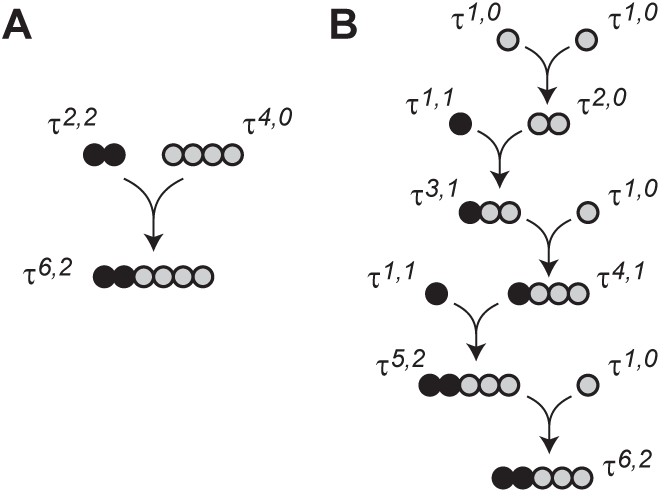
(A) Condensation between a pure ^13^C_2_ compound with MID *τ*^2,2^ and a C_4_ compound with natural distribution *τ*^4,0^ = *β*^4^, yielding the MID *τ*^6,2^. (B) Example of step-wise condensations of pure ^13^C atoms with natural ^13^C atoms, resulting in the same MID *τ*^6,2^.

It is straightforward to show that this convolution yields the MID

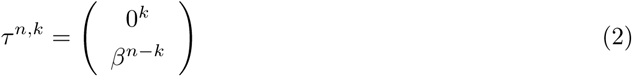

where 0^*k*^ denotes the *k*-dimensional zero vector. That is, convoluting a natural MID with the unit vector *e*^*k,k*^ will “shift” the MID by *k* steps. For example, the labeled compounds in Figure 2 would have MIDs

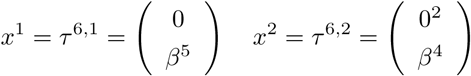

This model based on condensations reflects how larger molecules are synthesized by anabolic pathways, which is indeed how most tracers are produced: for example, the MID *τ*^6,2^ might be produced by a condensation *τ*^6,2^ = *τ*^2,2^ ⊗ *τ*^4,0^ (Figure 3A). Moreover, any sequence of condensations (in any order) that incorporate a total of *k* ^13^C atoms from a pure ^13^C source and *n* − *k* atoms from a natural carbon source will generate same MID *τ*^*n,k*^. This is because a condensation between any two *τ* distributions always yields another *τ* distribution,

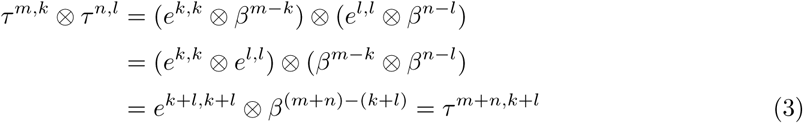

since the ⊗ operator is commutative and associative, and it holds that *β*^*m*^ ⊗ *β*^*n*^ = *β*^*m*+*n*^ for any *m, n*. Applying this property repeatedly, we see that an MID *τ*^*n,k*^ can be formed from 1-carbon building block as

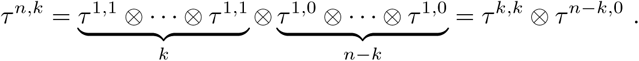

where the condensations can occur in any order. For example, Figure 3B shows one possible sequence of condensations giving rise to *τ*^6,2^ = *τ*^2,2^ ⊗ *τ*^4,0^.

With the class of distributions *τ*^*n,k*^, we can now model all MIDs in the example of Figure 2 as *x*^0^ = *τ*^6,0^, *x*^1^ = *τ*^6,1^ and *x*^2^ = *τ*^6,2^, and this generalizes to any mixture of compounds produced by condensations. However, the resulting mixture MID *x* in Figure 2 is not itself a *τ* distribution.

### 2.3 From mixture models to MID correction

We now derive the general case of the MID correcting transform *T*^*n*^ from the distributions *τ*^*n,k*^. Fernandez et al. [1996] described the case of an arbitrary *n*-carbon molecule formed as a mixture of *n* + 1 compounds with distributions *τ*^*n*,0^, τ^*n*,1^, …, *τ*^*n,n*^, defined as above. We can write the resulting mixture model in matrix form as

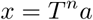

where *a* is again the vector of mixture coefficients, and *T*^*n*^ is the (*n* + 1) × (*n* + 1) matrix

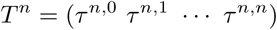

whose columns are the MID vectors of the mixed compounds. Comparing with (2), we see that this is exactly the MID correction matrix (1), if the vector of mixture coefficients *a* can be interpreted as the corrected MID *y*. To understand why this interpretation is valid, consider the following thought experiment (Figure 4).

For any metabolite with any MID *x*, we can partition the population of molecules into *n* + 1 subpopulations, where subpopulation *k* consists of those molecules that have incorporated *k*^13^C atoms from a pure ^13^C source and *n*−*k* atoms from a natural source, *k* = 0, 1, *…, n* (Figure 4A). Then, the MID of subpopulation *k* is precisely *τ*^*n,k*^, and *a*_*k*_ is the the fraction of molecules belonging to this sub-population, and we again have the mixture MID *x* = *T*^*n*^*a*. Now, imagine that we could replace all natural-derived ^13^C in these molecules with ^12^C (Figure 4B). Then the MID of subpopulation *k* would simply be the unit vector *e*^*n,k*^, and therefore the mixture MID becomes identical to the mixture coefficients, 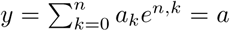. Therefore, the vector *a* can be interpreted as the MID *y* that we would have obtained if natural ^13^C did not exist, that is, the corrected MID.

**Figure 4:**
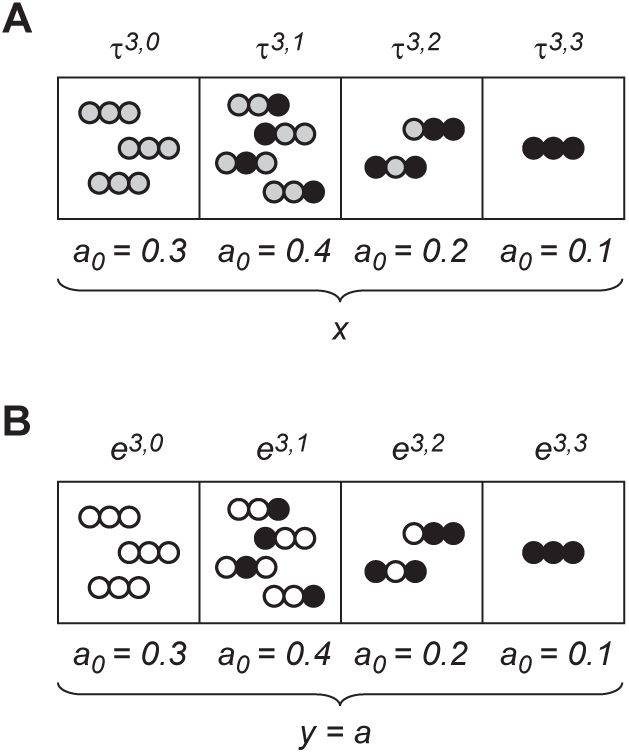
A thought experiment. (A) A pool of molecules partitioned into sets based on the number of ^13^C atoms derived from a pure ^13^C source (black). Gray indicates atoms drawn from the natural isotope distribution. (B) The same partition, with natural-distributed atoms replaced by ^12^C atoms (white). For details, see text.

In summary, we have demonstrated that for any *n*-carbon compound formed by combining combining pure ^13^C carbon with natural-distributed carbon, the observed MID *x* and the corrected MID *y* are related as *x* = *T*^*n*^*y*. This model is valid for any product of a metabolic network whose substrates are either pure ^13^C (*e*^*n,n*^) or natural (*β*^*n*^).

## 3 Properties of the transform *T*^*n*^

In this section, we discuss some interesting properties of the transform *T*^*n*^. To gain a bit of intuition, Figure 5 visualizes the effects of *T*^*n*^ for the case *n* = 2 at various points in the 2-simplex *S*_2_. (The *n*-simplex *S*_*n*_ is the set 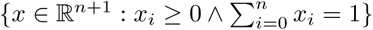, that is, the set of all possible *n*-carbon MIDs.) As with any linear operator, *T*^*n*^ maps the *k*’th unit vector *e*^*n,k*^ to the MID given by the *k*’th column of *T*^*n*^, *T*^*n*^*e*^*n,k*^ = *τ*^*n,k*^, *k* = 0, …, *n*. A special case is *k* = *n* for which *T*^*n*^*e*^*n,n*^ = *τ*^*n,n*^ = *e*^*n,n*^.

**Figure 5:**
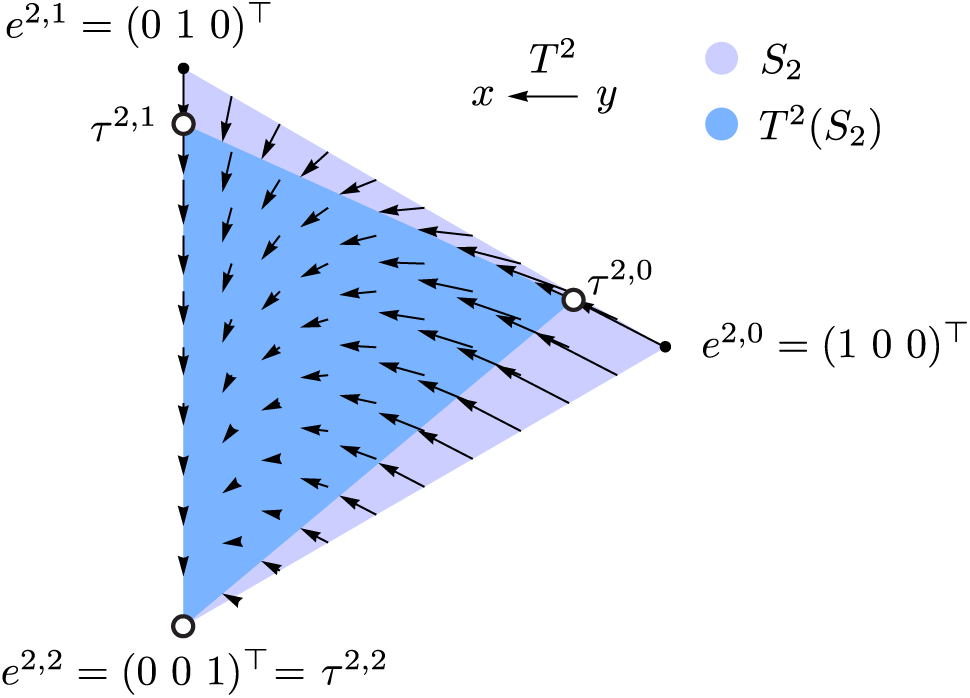
The effect of the transform *T*^2^ (for a 2-carbon compound) on MID vectors in the 2-simplex *S*_2_, drawn projected from R3 into the plane. Arrows indicate the effect of applying *T*^2^ at various points *y* ∈ *S*_2_. Dark blue region is the image *T*^2^(*S*_2_) of the 2-simplex, whose corners are the MIDs *τ*^2,*k*^, *k* = 0, 1, 2. Here the heavy isotope probability *p* was set to 0.1 for clarity.

This is the only eigenvector of *T*^*n*^ that lies within the simplex *S*_*n*_, reflecting the fact that a pure 13C_*n*_ compound cannot have incorporated natural ^13^C. All other vectors *y ∈ S_n_* are “pushed” towards *e*^*n,n*^ by *T*^*n*^ (Figure 5). Conversely, (*T*^*n*^)^−1^ maps each MID *τ*^*n,k*^ to the unit vector *e*^*n,k*^, and in particular maps the natural MID *τ*^*n*,0^ = *β*^*n*^ to the unit vector *e*^*n*,0^. It is in this sense that (*T*^*n*^)^−1^ “removes” natural ^13^C. However, it should be noted that there is nothing about the corrected MID *y* = (*T*^*n*^)^−1^*x* that distinguishes it as inherently free of natural ^13^C: we could in principle observe the same MID *y* from another measurement. This is evident from the fact that (*T*^*n*^)^−1^ is not a projection, as (*T*^*n*^)^−1^(*T*^*n*^)^−1^*x* ≠ (*T*^*n*^)^−1^*x*.

It is important to note that, while *T*^*n*^ is invertible, is it not surjective. It is clear that if *y* ∈ *S*_*n*_, then *x* = *T*^*n*^*y* is also in *S*_*n*_, since *T*^*n*^*y* is a convex combination of the columns of *T*^*n*^, all of which are in *S*_*n*_. But not all vectors *x* ∈ *S*_*n*_ can be written as *x* = *T*^*n*^*y*; that is, the image of *T*^*n*^(*S*_*n*_) does not cover all of *S*_*n*_ (Figure 5). For example, no unit vector *e*^*n,k*^ for *k* < *n* is in *T*^*n*^(*S*_*n*_). The part of *S*_*n*_ that lies outside *T*^*n*^(*S*_*n*_) are those MIDs cannot be corrected by (*T*^*n*^)^−1^. These are MIDs that cannot be formed by condensing pure ^13^C and natural-distributed carbon as described in Section 2.2. Generally, such MIDs vectors contain *less* ^13^C than expected from the natural distribution. MID vectors of this kind can be generated by measurement errors, for example if a mass isotopomer peak is underestimated in mass spectrometry analysis. Attempting to apply (*T*^*n*^)^−1^ to such an MID *x* will yield an *y* outside *S*_*m*_, and can result in negative elements of *y*, as other have observed [Millard et al., 2012].

Regarding numeric stability of the transform, it is easy to show that the eigenvalues of *T*^*n*^ are *λ*_*k*_ = (1 − *p*)^*k*^, *k* = 0, … *n*, which gives the matrix condition number *λ*_0_/*λ*_*n*_ = (1 − *p*)^−*n*^. Although this number increases exponentially with molecule size *n*, since 1−*p* is close to 1, it remains small for practically relevant values of *n*. For example, with *p* = 0.01, we have (1 − *p*)^−50^ = 1.65, indicating that *T*^*n*^ is stable for most practical purposes.

## 4 Flux analysis on corrected MIDs

We now turn to the question of how MID correction might affect metabolic flux analysis. We will restrict here ourselves to the case of metabolic and isotopic steady-state. A metabolic network acts on a set of substrates with MIDs **X**_in_ to generate products with MIDs **X**. Here we use bold **X** to indicate a set of MID vectors of various lengths. Each MID in **X**_in_ may be any *x* ∈ *T*^*n*^(*S*_*n*_), including ^13^C tracers and endogenous compounds. At steady-state, given a flux vector *v*, product MIDs **X** are uniquely determined by a nonlinear function **X** = *f*_*v*_(**X**_in_) of the substrate MIDs [Wiechert and de Graaf, 1997]. We will refer to *f*_*v*_ as the *network function*. Write **X** = **TY** to indicate the transform **T** that maps each *n*-carbon MID as *x* = *T*^*n*^*y*. For flux analysis on corrected MIDs to be valid, it must hold for any flux vector *v* that

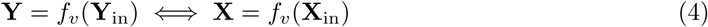

This criterion says that, if the observed product MIDs **X** are generated from the substrates **X**_in_ by the network, then the corrected product MIDs **Y** will be generated by the same network from the corrected substrate MIDs **Y**_in_, and vice versa. In other words, a flux vector *v* will fit the observed MIDs if and only if it fits the corrected MIDs, so metabolic flux analysis will give the same results in both cases. For (4) to hold true, it will be essential that we can exchange order between *f*_*v*_ and **T**. The following theorem is key.

### Theorem 1.

*For any metabolic network and flux vector v, the network function f*_*v*_ *satisfies*

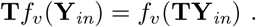

Theorem 1 implies the equivalence (4) because, if **Y** = *f*_*v*_(**Y**_in_), then **X** = **TY** = **T***f*_*v*_(**Y**_in_) = *f*_*v*_(**TY**_in_) = *f*_*v*_(**X**_in_). Conversely, if **X** = *f*_*v*_(**X**_in_), then **Y** = **T**^−1^*f*_*v*_(**X**_in_) = *f*_*v*_(**T**^−1^**X**_in_) = *f*_*v*_(**Y**_in_) as well. An illustration of the theorem for an example metabolic network is shown in Figure 6. We will first prove Theorem 1 for the special case of a compartmental network, and then prove the general case by representing a metabolic network as a cascade of compartmental networks connected by condensations, as described previously [Wiechert et al., 2001, Antoniewicz et al., 2007].

**Figure 6:**
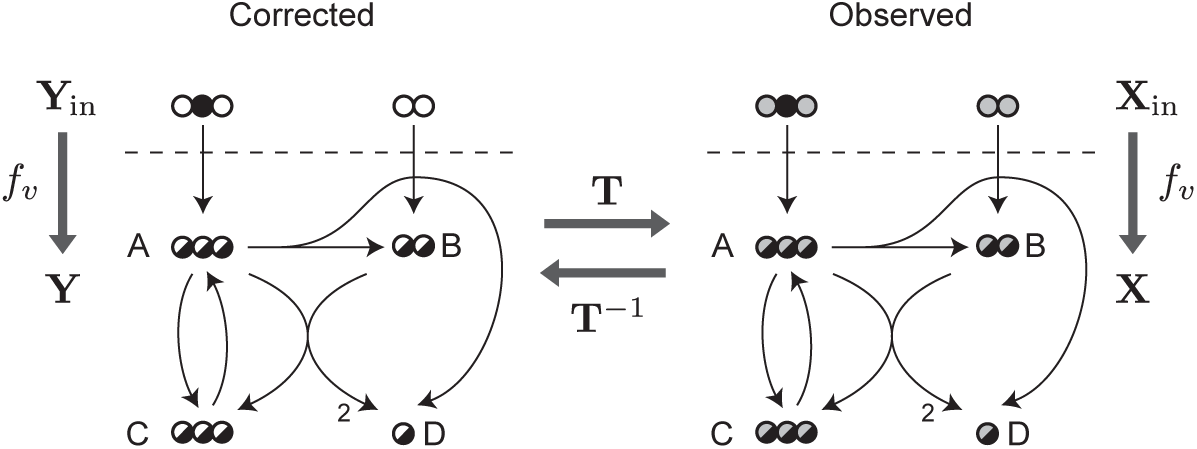
An illustration of Theorem 1 for an example metabolic network, adapted from [Antoniewicz et al., 2007]. Left, corrected mass isotopomers **Y**, where substrate atoms are either ^13^C (black) or ^12^C (white). Right, observed mass isotopomers **X**, where gray atoms indicate natural isotopic distribution. Half-black atoms indicate atoms whose isotopic state depends on the network function *f*_*v*_.

### 4.1 Compartmental networks

A compartmental network is a directed graph over *M* substrates and *N* products, all of which are *n*-carbon metabolites. At steady-state, for a given flux vector *v*, the compartmental network satisfies the mass balance equations

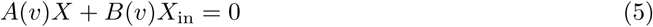

where *X* is an *N* × (*n* + 1) matrix whose rows are the product MIDs, and *X*_in_ is an *M* × (*n* + 1) matrix of substrate MIDs. The *N* × *N* matrix *A*(*v*) describes fluxes between products, and is always invertible [Anderson, 1979], while the *N* × *M* matrix *B*(*v*) describes fluxes from substrates to products. In this case, the network function *f*_*v*_ is found by solving the linear system (5), *X* = *f*_*v*_(*X*_in_) = −*A*(*v*)^−1^*B*(*v*)*X*_in_, and is easy to demonstrate that **T** and *f*_*v*_ can be exchanged.

#### Lemma 1.

*Let f*_*v*_ *be the network funcion of a compartmental network over n-carbon metabolites. Then it holds that*

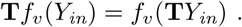

*Proof.* In this case, we can write the transform **T** as a matrix multiplication *X* = (*T*^*n*^*Y*^*T*^)^*T*^ = *Y* (*T*^*n*^)^*T*^. Since *T*^*n*^ and *f*_*v*_ are both linear, we obtain

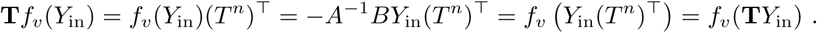

### 4.2 General metabolic networks

Any metabolic network can be written as a cascade of compartmental sub-networks, for molecule sizes *s* = 1, 2, …, *S*, connected by condensations that merge two smaller molecules into a larger one [Wiechert et al., 2001, Antoniewicz et al., 2007]. In this model, “molecules” need not be physical compounds, but can also represent various moieties of such compounds; however, this makes no difference for our purposes. At steady-state, each compartmental subnetwork satisfies the balance equation

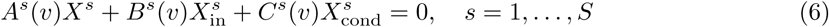

where *A*^*s*^(*v*), *B*^*s*^(*v*) and *C*^*s*^(*v*) are matrices describing the subnetwork structure, *X*^*s*^ and 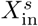 product and substrate MIDs matrices as before, and 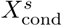 is an additional MID matrix formed by condensations *x*^*s*^ = *x*^*m*^ ⊗ *x*^*n*^, where *x*^*m*^ and *x*^*n*^ are MIDs from *X*^*m*^ and *X*^*n*^, for some *m, n* such that *m* + *n* = *s*. Thus, each 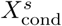 is a function 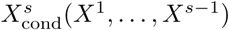 of the product MIDs from smaller size subnetworks. Let 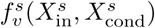 denote the network function for compartmental subnetwork *s*. Viewing 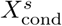 as a known constant, 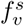is again a linear function, given by solving (6). The entire metabolic network then has a network function 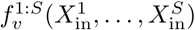 defined as

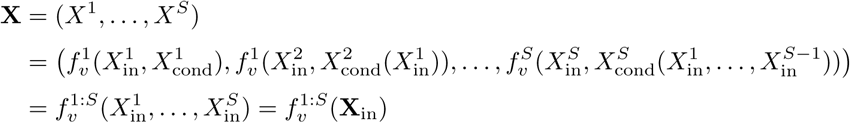

Because each 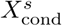 is a formed by convolution of the products from smaller size subnetworks, the proof of Theorem 1 will require that we also can exchange the convolution operator ⊗ with the transform **T**, so that 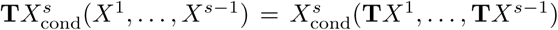. This is guaranteed by the following lemma.

#### Lemma 2.

*For any two MID vectors x*^*m*^ ∈ *S*_*m*_ and *x*^*n*^ ∈ *S*_*n*_, *it holds that*

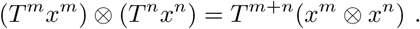

A proof is given in Appendix A. We are now ready to prove Theorem 1 for the general case.

*Proof of Theorem 1.* We will proceed by induction over the size *t* of the largest subnetwork. For *t* = 1 the theorem is true by Lemma 1. For *t* > 1, we assume that the theorem holds for the network formed by subnetworks *s* = 1, …, *t* − 1, that is, 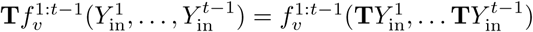. For the size *t* compartmental subnetwork, we have

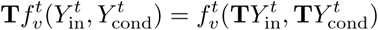

by Lemma 1, and

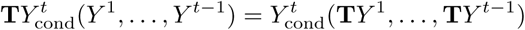

by Lemma 2. Together with the induction assumption, this gives

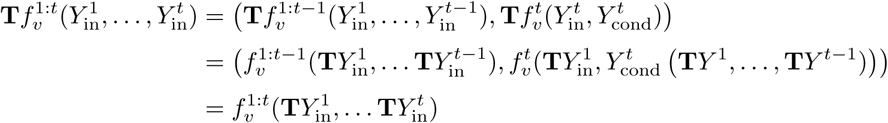

which completes the induction step.

## 5 Discussion

In this paper, we have provided a self-contained derivation of the linear transform *T*^*n*^ commonly used for isotope correction, and explored the effects of *T*^*n*^ for metabolic flux analysis. We hope that this contribution will help clarify the role of isotope correction in analysis of mass spectrometry data from ^13^C tracing experiments. It should be clear from our derivation that *T*^*n*^ is not an empirical error correction (as the name might suggest), but a rather a transformation that yields a particularly simple representation, the theoretical “corrected” MID *y*, which by definition contains no natural-derived ^13^C. At first glance, it might seem remarkable that a single linear transform exists that can “remove” ^13^C from any observed metabolite, with any isotopomer distribution whatsoever. This is possible because we have assumed a particular underlying model generating labeled compounds, namely that ^13^C enters metabolites by condensations of pure ^13^C atoms with natural ^13^C distributed atoms (section 2.2). This process always yields MIDs of the type *τ*^*n,k*^ for an *n*-carbon compound with *k* heavy atoms, and the argument in section 2.3 shows that (*T*^*n*^)^−1^ maps any mixture of *τ*^*n,k*^ MIDs to a corrected MID *y*. However, it is important to note that (*T*^*n*^)^−1^ is not applicable to MID vectors that cannot be generated by the process we assumed; in particular, MIDs with less ^13^C enrichment than the natural distribution are not feasible. This can cause problems in cases where the measured mass isotopomer fractions are biased or uncertain.

In our derivation of the class of distributions *τ*^*n,k*^, we only considered the case of pure ^13^C tracers. In practise, tracers always have some degree of ^12^C impurities. To handle this complication, the 13C source can be considered to emit ^13^C with a propability *q* < 1 (where *q* is typically around 0.98 − 0.99), and the pure ^13^C distribution *e*^*k,k*^ is then replaced with another binomial, say *α*^*n*^ = Bin(*n, q*). We would then obtain a different labeled compound distribution *τ*^*n,k*^ = *α*^*l*^ ⊗ *β*^*n*−*k*^, and can construct a new linear transform *T*^*n*^ based on this distribution. Importantly, equation (3) still holds for such *τ* distributions, and therefore the proof of Lemma 2 remains valid. Hence, our main result is still applicable in this more general case.

One practical advantage of isotope correction is that the vectors *y* are typically sparse, since only atoms that derive from the pure ^13^C source (that is, the tracer) can yield nonzeros in *y*. Hence, in the typical isotope tracing experiment, most mass isotopomer fractions *y*_*i*_ will be zero. This sparsity is intuitively exploited by researchers creating metabolic network models, by manually excluding moieties that cannot possibly be reached by tracer-derived atoms, such as cofactors and essential amino acids. However, this process could be performed automatically for any given metabolic network, by discarding any atoms that cannot be reached from tracer ^13^C. For example, in a previously published model of one-carbon metabolism with U-^13^C_5_-methionine as tracer [Nilsson et al., 2017], a total of 341 atoms had to be included to simulate the measured MIDs, but only 65 atoms are needed to simulate the corresponding corrected MIDs.

Although it is clear that the isotope-correcting transform is widely applicable, it is not obvious that metabolic flux analysis (MFA) on a set of corrected MIDs **Y** is equivalent to MFA on the corresponding set of observed MIDs **X**. Our theorem 1 provides a theoretical foundation for using MFA on corrected MIDs, by guaranteeing that any set of steady-state isotopomer balances (6) hold for **Y** if and only if they hold for **X**. In other words, isotopomer balances for **Y** and **X** are compatible with precisely the same set of flux vectors *v*. This equivalence holds because *T*^*n*^ is linear and commutes convolution of MIDs (Lemma 2). However, an important limitation is that Theorem 1 applies only when the isotopomer balances hold exactly. In practice, where models are inaccurate and observed MIDs **X** are afflicted by measurement errors, there is generally no vector *v* that that exactly satisfies **X** = *f*_*v*_(**X**_in_). Instead, we must seek a *v* that minimizes the error ||**X** = *f*_*v*_(**X**_in_)||, where ||⋅|| typically is the *L*_2_ (Euclidean) norm. This will in general *not* yield the same solution as minimizing ||**Y** = *f*_*v*_(**Y**_in_)||, since the *L*_2_ norm will measure errors differently in the two representations. Consequently, both point estimates and confidence intervals on fluxes will generally differ between the two representations. This issue should be further investigated, and we would recommend that flux solutions from MFA on corrected data **Y** should be confirmed to also fit the uncorrected data **X**. Interestingly, this problem could also be viewed as an indication that the *L*_2_ norm is not appropriate for this setting, and it could be argued that a suitable norm should generate identical solutions in both representations. Whether there exists a norm on the *n*-simplex *S*_*n*_ satisfying this criterion, for example one as the measures described by Aitchison [1982], is an interesting topic for future studies. Finally, we have only treated metabolic networks at metabolic and isotopic steady-state. It seems that the proof strategy employed here should also extend to isotopic non-steady state (which involves a cascade of differential equations of a similar structure), but a rigorous proof for this scenario clearly requires further work.

## Acknowledgements

The author wishes to thank Dr. Firas Midani for helpful suggestions on the manuscript. This work was funded by the Foundation for Strategic Research grant no.ITM17-0245 and Karolinska Institutet.

## A Proof of Lemma 2

*Proof of Lemma 2.* For the case of basis vectors *x* = *e*^*m,k*^ and *y* = *e*^*n,l*^, we already know that

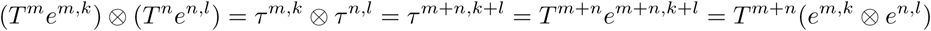

from equation (3). For arbitrary vectors *x* ∈ *S*_*m*_ and *y* ∈ *S*_*n*_, write 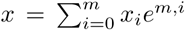 and 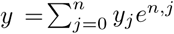. Then

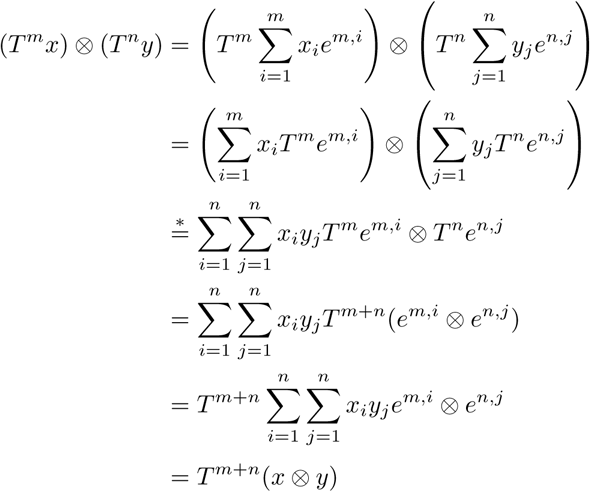

where in step ∗ we have used the fact that the ⊗ operation distributes over vector addition, that is, *a* ⊗ (*b* + *c*) = *a* ⊗ *b* + *a* ⊗ *c* for any vectors *a, b, c*.□

## Notes

### Competing Interest Statement

The authors have declared no competing interest.

